# search.bioPreprint: a discovery tool for cutting edge, preprint biomedical research articles

**DOI:** 10.1101/052563

**Authors:** Carrie L. Iwema, John LaDue, Angela Zack, Ansuman Chattopadhyay

## Abstract

The time it takes for a completed manuscript to be published traditionally can be extremely lengthy. Article publication delay, which occurs in part due to constraints associated with peer review, can prevent the timely dissemination of critical and actionable data associated with new information on rare diseases or developing health concerns such as Zika virus. Preprint servers are open access online repositories housing preprint research articles that enable authors (1) to make their research immediately and freely available and (2) to receive commentary and peer review prior to journal submission. There is a growing movement of preprint advocates aiming to change the current journal publication and peer review system, proposing that preprints catalyze biomedical discovery, support career advancement, and improve scientific communication. While the number of articles submitted to and hosted by preprint servers are gradually increasing, there has been no simple way to identify biomedical research published in a preprint format, as they are not typically indexed and are only discoverable by directly searching the specific preprint server websites. To address this issue, we created a search engine that quickly compiles preprints from disparate host repositories and provides a one-stop search solution. Additionally, we developed a web application that bolsters the discovery of preprints by enabling each and every word or phrase appearing on any website to be integrated with articles from preprint servers. The bioPreprint search engine is publicly available at http://www.hsls.pitt.edu/resources/preprint.

## Introduction

Preprint servers are online repositories that manage access to manuscripts that have not yet been peer-reviewed or formally published in a traditional manner. Preprint manuscripts are not copyedited, but they do undergo a basic screening process to check against plagiarism, offensiveness, and non-scientific content. Authors may make revisions at any point, but all versions remain available online. It should be noted that the term “preprint” in this context refers to manuscripts posted by the authors themselves onto specific online servers, not articles made available online by publishers a few weeks ahead of traditional publication

Preprint articles can be more difficult to discover than those published traditionally, as they are not currently indexed in Medline and therefore do not appear in PubMed search results. This suggests that many timely and relevant research reports potentially fall through the cracks, as the time it takes to traditionally publish a biomedical manuscript can take anywhere from a few months to a few years. This lengthy process is seen by researchers to be a hindrance to scientific advancement. In response, there is a developing movement of preprint advocates who propose that preprints play a role in “catalyzing scientific discovery, facilitating career advancement, and improving the culture of communication within the biology community” (1). Preprint servers “enable authors to make their findings immediately available to the scientific community and receive feedback on draft manuscripts before they are submitted to journals” (2).

The history, rationale, and controversy surrounding preprint servers and the pace of the current publication process has been well addressed in other manuscripts (3–14), news items (15–22), and blogs or white papers (23–32). We do not intend to duplicate this information here, but suggest exploration of our reference list for an overview of the current state of the topic.

### Preprint Server Examples

There are currently only a small number of preprint servers catering to biological and biomedical research manuscripts.

- **<u>arXiv</u>** is a venerable preprint server covering physics, mathematics, computer science, nonlinear sciences, statistics, and quantitative biology since 1991. arXiv is funded by Cornell University Library, the Simons Foundation, and many member institutions.
- **<u>bioRxiv</u>**, operated by Cold Spring Harbor Laboratory, covers new, confirmatory, and contradictory results in research ranging from Animal Behavior and Cognition to Clinical Trials, Neuroscience to Zoology.
- **<u>F1000Research</u>**, a member of the Science Navigation Group, provides an open science platform for the immediate publication of scientific communication. Posters and slides receive a digital object identifier and are instantly citable. Articles with associated source data are published within a week and made available for open peer review and user commenting. Articles that pass peer review are then indexed in PubMed, Scopus, and Google Scholar.
- **<u>PeerJ Preprints</u>** covers biological, medical, life, and computer sciences. Their aim is to reduce publishing costs while still efficiently publishing innovative research, with an emphasis on not yet peer-reviewed articles, abstracts, or posters. Submissions are free, can be a draft, incomplete, or final version, and are typically online within a day after editorial approval.

Our intention is to present a resource that facilitates the quick and easy identification and access of scientific content located on preprint servers. <u>The Health Sciences Library System at the University of Pittsburgh</u> (HSLS) developed a tool to help researchers to quickly search preprint databases and discover cutting edge, yet-to-be published or reviewed biomedical research articles, <u>search.bioPreprint</u> (**Figure 1**). This search engine encompasses a federated search of arXiv, bioRxiv, F1000Research, and PeerJ Preprints. We chose to publish this article on a preprint server in order to support the preprint movement and to elicit feedback on usage of the tool, which will be updated as needed.

**Figure 1:**
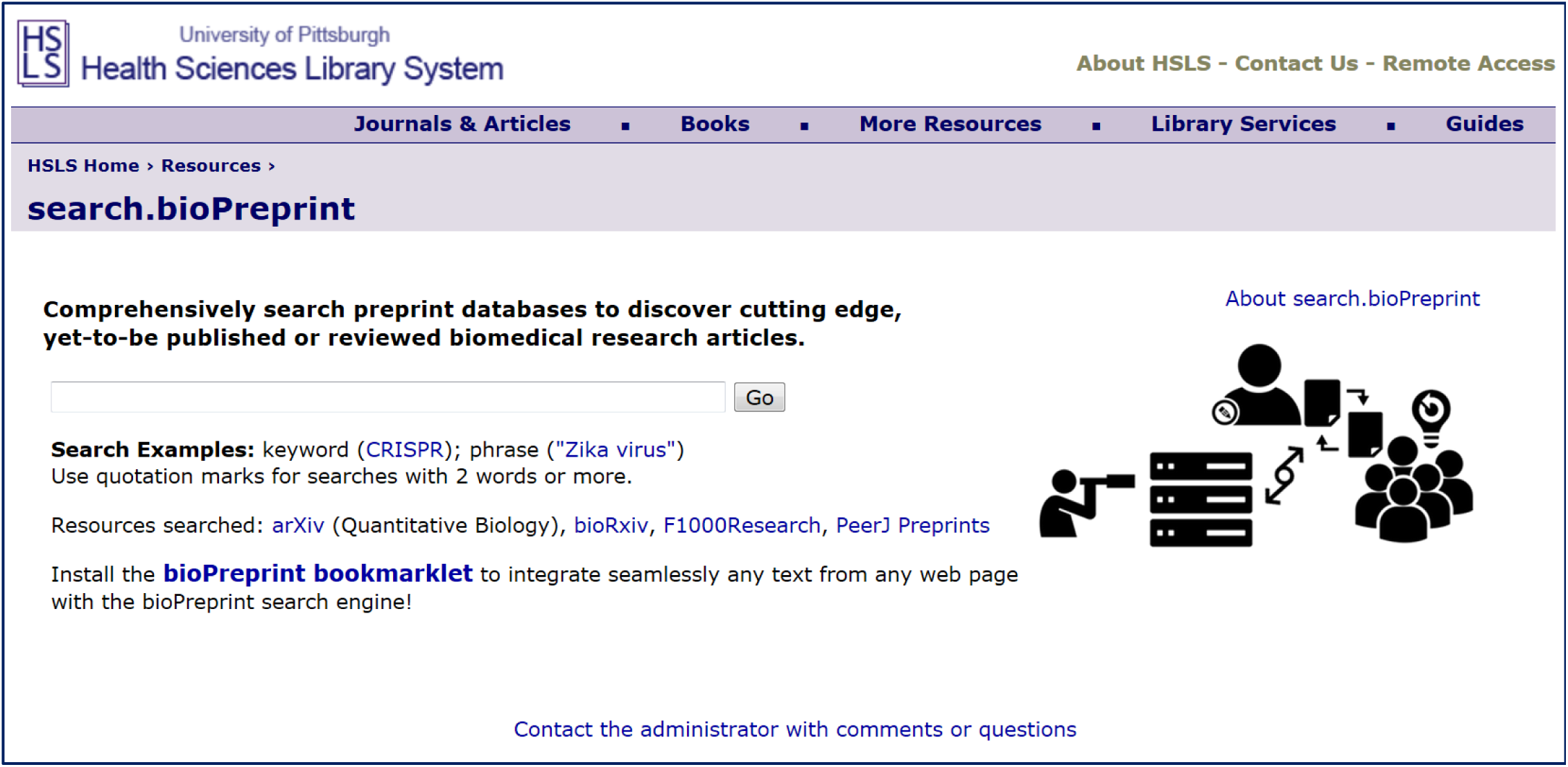
Website homepage for <u>search.bioPreprint</u>.

## Implementation

### Search Engine

Search.bioPreprint was created using <u>IBM Watson Explorer</u> to generate a meta search engine that compiles search term results from a pre-selected list of multiple sources into a single list ordered by the relevance of matching query terms. The results can then be further filtered by Source (e.g., the preprint servers of origin) or by Topic (e.g., microcephaly for a Zika Virus search). The Topic search is accomplished via clustering, meaning the search results are organized on the fly by similarity in subject matter. Additionally, a “remix” link displayed next to the clustered topics reveals new secondary topics. This is done by clustering the same search results again, but explicitly ignoring the topics that were used in the initial clustering process.

The Health Sciences Library System at the University of Pittsburgh has repeatedly utilized IBM Watson Explorer (formerly Vivisimo Velocity) software to develop, implement, and maintain several federated search engines focused on a variety of topics. These include: <u>search.HSLS.OBRC</u>–a portal for discovering bioinformatics databases and software via the Online Bioinformatics Resource Collection (33), <u>Clinical Focus</u>–a portal providing quick access to high-quality clinical information (34), and <u>Clinical eCompanion</u>–a portal with information for primary care (35).

The selected sources for retrieving preprint articles are: (1) the quantitative biology section of <u>arXiv.org</u>, (2) <u>bioRxiv</u>, the preprint server for biology, (3) <u>F1000Research</u> and (4) <u>PeerJ Preprints</u>. A maximum of 200 total results are returned based on the licensing agreement with IBM Watson Explorer; this also contributes to a short wait for return of results.

### How-To-Use

As an example, typing a single-word query term, such as CRISPR, into the search box results in <u>ninety-one preprint articles</u> culled from the aformentioned preprint servers (**Figure 2**). Clicking on an article title redirects to that article at its original source. Search results may be narrowed by Topic or Source using the filters on the left side of the page. Using the CRISPR example, the ninety-one search results are grouped into shared Topics: fourteen articles on “Bacterial,” twelve articles on “Protein,” six articles on “Genome engineering,” etc. Expanding individual topics reveals a list of subtopics: clicking on the topic “Protein” redistributes the twelve articles into subtopics, including “CRISPR-Cas9,” “Image, Palindromic Repeat,” “Mutants, Generated,” etc. Clicking on a topic or subtopic reconfigures the search results to limit to these filtered articles.

**Figure 2:**
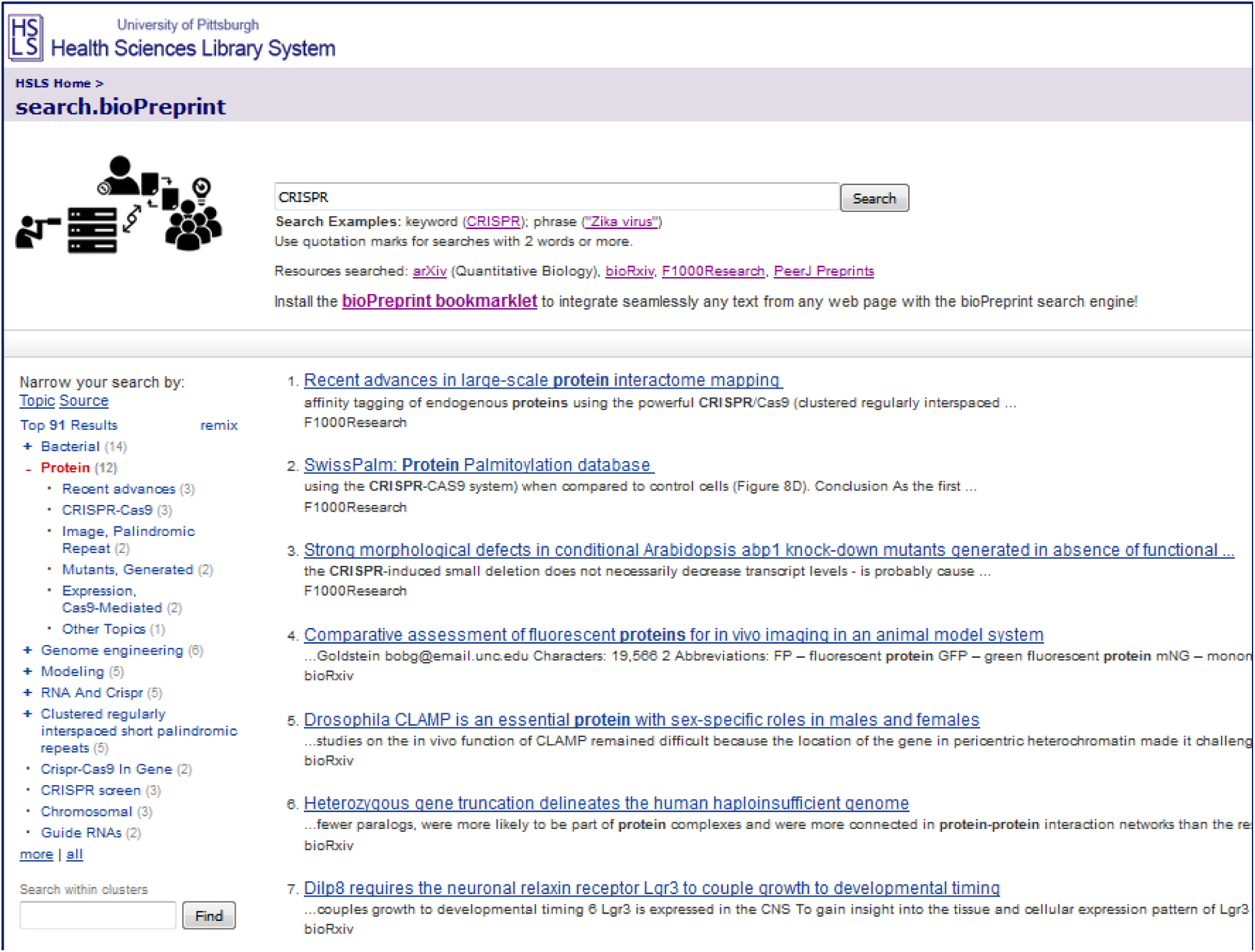
Search results page with query term CRISPR. At left is the default view by Topic. (2 May 2016)

Clicking on the “remix” button appearing next to “Top 91 results” regroups the original search results into additional topics such as “Cells,” “Advances,” “Drosophila,” etc that are not present in the first results iteration (**Figure 3**). This provides another opportunity to discover pertinent preprint articles, especially if a large number of results is returned.

**Figure 3:**
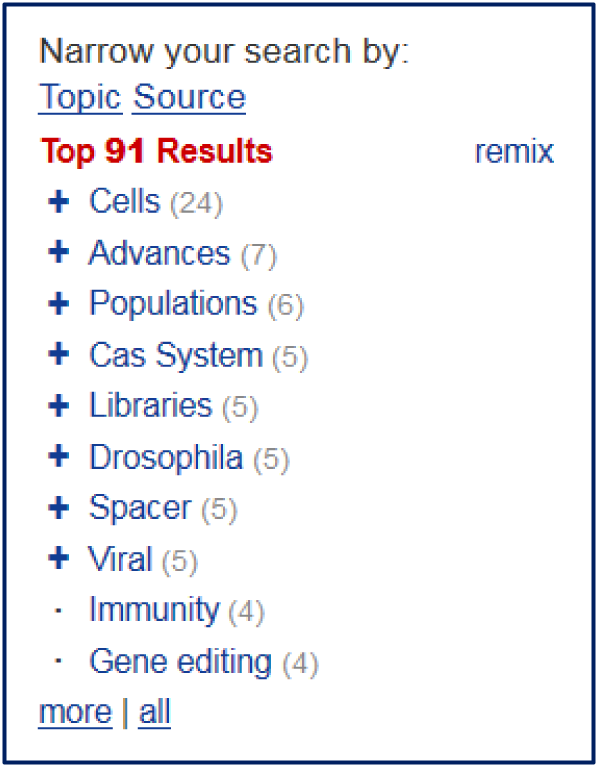
Topics change after selecting “remix.” (2 May 2016)

The search results may also be filtered by Source. Selecting this will change the default display of Topic-focused clusters to articles organized by Source, which in the current iteration is one of the four preprint servers searched by this tool: nineteen from F1000Research, two from PeerJ Preprints, six from arXiv, and sixty-five from bioRxiv (**Figure 4**).

**Figure 4:**
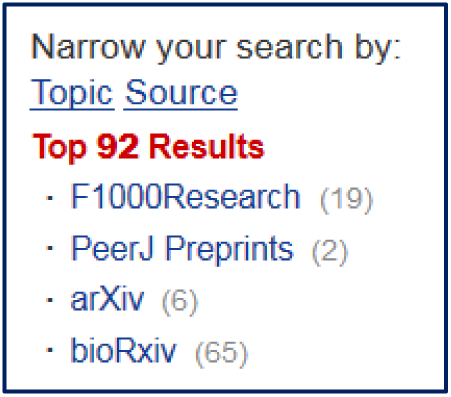
Results view by preprint server Source. (2 May 2016)

Quotation marks are recommended for searches with exact phrases, e.g., Zika virus. The necessity of this was discovered after examing the search parameters of the various preprint servers. As one of the preprint servers by default joins words in a multi-word query with the Boolean operator “OR” then a search for a phrase such as zika virus produces multiple articles where the only matching term is virus. Using quotation marks for a search of more than one word mitigates this problem and considerably improves the quality of results. A search for “zika virus” thus produces <u>seventy-nine articles</u> that are topically filtered into “Zika virus infection,” “Microcephaly,” “Discovery,” “Dengue Virus,” etc.

The “Search within clusters” box allows for searching within the search results, and can be used to identify specific articles within the cohort of Zika virus preprints that are not immediately apparent from Topical clustering. Entering vaccine in the search box highlights the topics and subtopics containing articles bearing the word vaccine: under “Zika virus infection” is “Preventing Zika Virus Infection;” under Dengue Virus is “Antibodies, Vaccine” and “Community, Vector.” Selection of highlighted topics or subtopics reconfigures the results to limit to vaccine-related Zika virus preprints (**Figure 5**).

**Figure 5:**
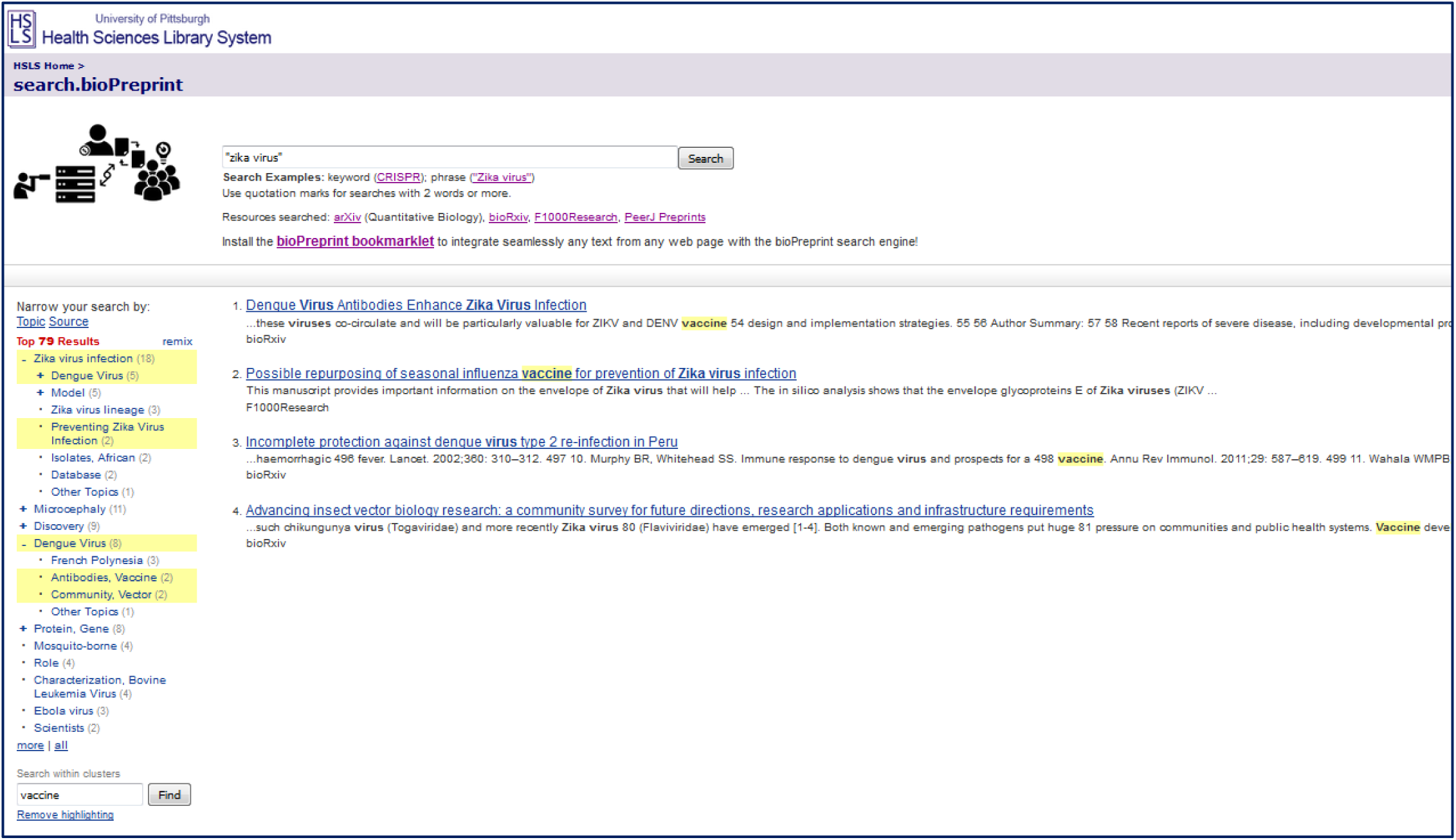
Results for Zika virus using quotation marks and the “Search within clusters” feature. (2 May 2016)

### bioPreprint-Bookmarklet

A bookmarklet is a special type of web browser widget containing an embedded software command, typically written in JavaScript, that extends the application of the browser by adding a one-click function as a bookmark. We created a <u>bioPreprint-bookmarklet</u> in order to seamlessly integrate a search for any word or phrase from any web page with the information stored in preprint servers. After dragging/dropping the bioPreprint-bookmarklet into any web browser, the next step is to highlight a word or phrase of interest then click the bookmarklet. This will result in a pop-up window displaying preprint articles containing the text of interest (**Figure 6**).

**Figure 6:**
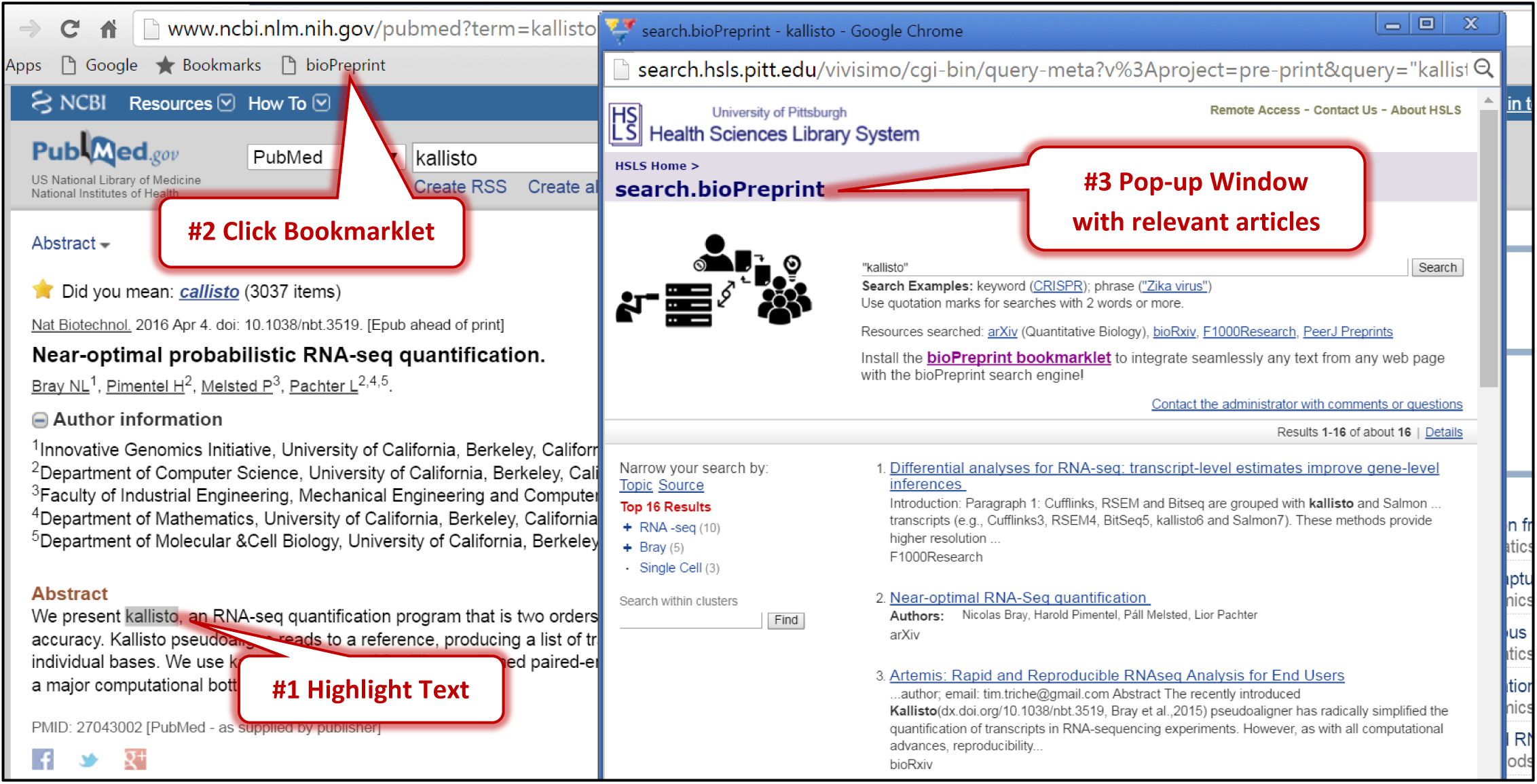
Using the bioPreprint-bookmarklet. (2 May 2016)

All web browsers that support JavaScript (Google Chrome, Mozilla FireFox, Internet Explorer, Apple Safari, Opera) are compatible with the bookmarklet. In case the favorites/bookmark bar is not visible we provide <u>instructions</u> for displaying it on commonly used browsers. A <u>video</u> describing how to install the bookmarklet in a web browser is also available.

## Use Cases

### Scenario 1

Imagine a researcher is searching PubMed for articles on “RNA-seq quantification” and comes across a paper recently published in Nature Biotechnology, “Near-optimal probabilistic RNA-seq quantification” (36). This paper introduces a new software program, Kallisto, that analyzes RNA-seq data by two orders of magnitude faster than previously used software. This is notable as it removes the computational bottleneck for RNA-seq data analysis. After reading about this new software, the researcher decides to check whether its been widedly adopted by perusing the published literature.

A search in PubMed with the search term “Kallisto” results in only the original article. This is well within expectations, considering the recent publication date of the article, 4 April 2016. There has not been enough time for researchers to know about the software, let alone write papers citing it.

To continue to try and gauge the usage of Kallisto in RNA-seq data analysis, the researcher might take an alternative approach: instead of searching PubMed, try searching for preprint articles. This can be achieved with a single click of the bioPreprint-bookmarklet once it is installed in the researcher’s web browser. Upon viewing the article abstract on the PubMed search results page, highlighting the word “Kallisto,” and clicking the bioPreprint-bookmarklet, a pop-up appears with the search.bioPreprint search results: sixteen preprint articles, two from arXiv, thirteen from bioRxiv, and one from F1000Research. Interestingly, the second article on the results page is the preprint version of the Nature Biotechnology paper on Kallisto software, submitted to the arXiv preprint server (**Figure 6**). The authors submitted their preprint on 11 May 2015, almost one year before its publication in Nature Biotechnology, with concomitant indexing by PubMed.

It is worth noting that since the availability of the Kallisto paper as a preprint, fifteen preprints have cited the use of Kallisto software. These articles cover numerous topics, including development of new software, single cell RNA-seq analysis, and quantification of the relative abundance of transcripts in various experimental settings.

### Scenario 2

A student gathering information from the internet about the regulation of gene expression happens upon the GTEx Project Community Scientific Meeting <u>website</u>. GTEx stands for the Genotype-Tissue Expression project (GTEx), which aims to develop an atlas of human gene expression and its regulation across various tissue types. Intrigued by the scope of this project, the student is curious to know how GTEx project data have been utilized in research.

The bioPreprint search engine and bookmarklet can quickly satisfy the student’s curiosity by providing easy access to GTEx-related articles hosted by various preprint servers that may or may not be published “in print” yet. This process is simple, unique, and the student doesn’t even need to leave the current web page to go on a literature hunt. Rather, all GTEx-related articles will appear in a new window with only two clicks, the first highlighting the word GTEx and the second on the previously-installed bioPreprint-bookmarklet. The result is sixty-seven articles showcasing the use of GTEx data in a variety of research topics including “Genome Wide Association Studies,” “Allele, Specific expression,” “Expression Quantitative Trait Loci,” etc.

## Conclusions

These use cases emphasize the power of the bioPreprint search engine and associated bookmarklet in delivering scientific research articles that are not only hard-to-find and yet-to-be traditionally published, but also on demand at the point of reading. And the “point of reading” can be anything on the web: journal articles, news items, blogs, PubMed/Google Scholar search results, etc.

Until the creation of search.bioPreprint there has been no simple way to identify biomedical research published in a preprint format, as they are not typically indexed and are only discoverable by directly searching the preprint server websites. search.bioPreprint is a one-stop-shop for finding these types of articles, and an important contribution to the preprint movement. During the final stages of manuscript preparation an online database aiming to index preprint articles was launched, <u>PrePubMed</u>, which despite appearances is not an official resource from the National Library of Medicine (NLM), the National Center for Biotechnology Information (NCBI), or PubMed. We want to acknowledge this new resource, but emphasize that search.bioPreprint offers not only full text searching, but also topical and source-based clustering of results. In addition, our tool has been available since mid-February 2016, around the same time as the <u>ASAPbio meeting</u>, where it was mentioned during discussions.

The underlying technology upon which search.bioPreprint was built is flexible enough to integrate additional resources into the search engine. As new preprint servers are introduced, search.bioPreprint will incorporate them and continue to provide a one-stop solution for finding preprint articles. We welcome feedback that introduces new preprint resources and addresses usability concerns.

The bioPreprint-bookmarklet enables each and every word or phrase appearing on any website to be integrated with information in articles stored in preprint servers. The on-demand delivery of preprint articles at the point of reading enables researchers to discover brand new pre-published articles quickly and be updated with cutting edge, yet-to-be-reviewed information that is challenging to discover by traditional literature searching methods. Our intention is that the combined use of the aforementioned tools helps to fulfill the unmet need of the scientific community for immediate dissemination of research outcomes, ultimately resulting in improved scientific communication and far-ranging insights and innovations.

## Limitations

The quality of the search results generated by the bioPreprint search engine is confined by the search parameters of the individual preprint servers. If the preprint servers alter their search algorithms, a concomitant adjustment of underlying codes used by the bioPreprint search engine is often required. Unfortunately, this can be done without any public notification and is only discoverable upon a thorough analysis of bioPreprint search results. The University of Pittsburgh Health Sciences Library System has a quality check team involving two librarians to ensure the accuracy of search.bioPreprint results. The team routinely compares the search results produced by several preset query terms with the previous results and reports any discrepancies to the development team.

The average time taken to display search results is not always optimal. The speed of the search.bioPreprint results return stems from multiple factors: individual preprint servers’ searching speed, efficiency of the IBM Watson Explorer software, and computational power of the server hosting the bioPreprint search engine. While some contributing factors are outside of our control, efforts will be undertaken to speed up the search process by continually upgrading the power of the host server.

## Data and Software Availability

- Search.bioPreprint is freely available at http://www.hsls.pitt.edu/resources/preprint.
- The bioPreprint-bookmarklet is freely available at http://hsls.pitt.edu/biopreprint-infobooster.
- The JavaScript code embedded in the bookmarklet is:
“javascript:(function(){(function(t,u,w){t=“+(window.getSelection%3Fwindow.getSelection():document.getS election%3Fdocument.getSelection():document.selection%3Fdocument.selection.createRange().text:”);u=t %3F‘http://search.hsls.pitt.edu/vivisimo/cgi-bin/query-meta%3Fv%253Aproject=pre-print%26query=%2522’+encodeURIComponent(t)+‘%2522’:“;w=window.open(u,‘_blank’,‘height=750,width=700,scrollbars=1’);w.focus%26%26
w.focus();if(!t){w.document.write(‘<html><head><title></title></head><body style=wpadding:1em;font-family:Helvetica,Arial”><br /><p>First%2C highlight a word or a group of words from any website that you are browsing (journal article%2C PubMed search result%2C news article%2C blog%2C etc.)%2C and then click on this bookmarklet to retrieve cutting edge%2C yet-to-be published or reviewed biomedical research articles related to your selected word(s).</p></a><p>Check the <a</p>
href=\“http://media.hsls.pitt.edu/media/BioPreprint_ac0316.mp4\”>How to Video</a> for instruction.</p><br /><p><img src=“http://www.hsls.pitt.edu/sites/all/themes/liberry_front/logo.png</underline>;” alt=“HSLS Logo”></p><script>var q=document.getElementById(“q”),v=q.value;q.focus();q.value=“”;q.value=v;</script></body></html>’); w.document.close();}})()})();”

## Author Contributions

AC conceived the concept and wrote the Implementation section. CI created the search.bioPreprint logo, assisted with concept refinement and design, and wrote all documentation, including preparation of the initial drafts of the manuscript. AC and CI created the figures. JL developed the search engine. AZ created the bookmarklet and webpage. All authors were involved in revision of the draft manuscript and have agreed to the final content.

## Competing Interests

The authors declare no competing interests.

## Grant Information

The authors declare that no grants were involved in supporting this work.

## Acknowledgements

### Acknowledgements

The authors wish to gratefully acknowledge the following individuals for their help with various aspects of the creation of search.bioPreprint and manuscript preparation: Peter Coles for writing a <u>blog</u> on the insightful use of bookmarklets, Julia Dahm for the creation of the video describing how to install the bookmarklet, Melissa Ratajeski for providing helpful comments on the manuscript, Nancy Tannery for providing helpful comments on the manuscript and offering general support for this project, and Fran Yarger for offering general support for this project.

### Supplementary Material

There is no supplementary material for this manuscript.

